# Male and female *Drosophila suzukii* maintain extended, stable flight headings to a discrete sun stimulus

**DOI:** 10.64898/2026.07.06.736788

**Authors:** Kei Horikawa, Konstantin Savkin, Logan Rower, Liam Hodge, Timothy L. Warren

## Abstract

Long-distance movement in insects has crucial impacts on agriculture, human health, and biodiversity. Although it was long assumed that only large, specialist insects had the navigation capacity to support long-distance dispersal, recent studies have demonstrated that smaller insects, such as the tiny fruit fly *Drosophila melanogaster*, can maintain extended, straight paths while flying or walking. This raises the question of whether other *Drosophila* species possess the navigation capacity to support extended dispersal. Resolving this question is particularly important for *Drosophila suzukii* (spotted-wing drosophila), a potent pest species that causes enormous damage worldwide to ripe fruit and berries. Spotted-wing drosophila has been thought to lack a capacity for long-distance dispersal, as prior studies have estimated maximal daily dispersal distances of less than 90 m. We developed a system to continuously track the flight trajectories of magnetically tethered *D. suzukii* relative to a discrete, overhead LED that mimicked the sun. We found that flies maintained remarkably straight flight headings that varied unpredictably across individuals. Male and female *D. suzukii* exhibited a similar navigation capacity; both sexes responded to rotation of a discrete sun stimulus with compensatory turns to maintain a stable relative heading. Our results suggest that *D. suzukii* has an underappreciated capacity for rapid, radial dispersal, which could exceed 250 m in 15 min. This capacity may contribute to the pest species’ invasiveness and its reliable, annual re-establishment in seasonally intolerable climates. Our findings highlight the importance of developing area-wide, regional strategies to manage dipteran pest species such as *D. suzukii*.

## Introduction

Insect movement at a landscape scale involves enormous shifts in biomass, with both beneficial and deleterious impacts on agricultural crops (Asplen et al., 2015; Klein et al., 2006). Moreover, long-distance dispersal helps maintain genetic diversity of insect populations and enables their survival in severe, changing climates (Nichols and Hewitt, 1994; Asplen, 2018). Certain large, specialist insects – including specific moths, dragonflies, butterflies, and beetles – have been long recognized for their capacity to migrate long distances at a stable heading. For instance, monarch butterflies and common green darner dragonflies can travel 143 km (> 4 million body lengths) and 122 km (∼1.5 million body lengths) in a day (Knight et al., 2019). Death’s-head hawkmoths can fly nearly 90 km (∼1.5 million body lengths) over a single night (Menz et al., 2022). Displacement at this scale and speed requires keeping a nearly straight course (Chapman et al., 2015; Warren et al., 2019; Menz et al., 2022). For flying insects, a common, robust strategy for traveling in a stable direction is to maintain a constant heading vector relative to stable external reference cues such as the sun, the moon, the pattern of polarized skylight, or distant landmarks (Chapman et al., 2015; el Jundi et al., 2014; Warren et al., 2019; Green et al., 2019; Weisman et al., 2025).

Although it is sometimes assumed that only large, specialist insects possess the neural machinery to navigate long distances, several recent studies have shown that the tiny, common fruit fly, *Drosophila melanogaster*, can maintain extended, straight trajectories using external reference cues. Tethered female *D. melanogaster* maintain stable flight headings to natural and simulated patterns of polarized skylight, and to point light sources mimicking the sun (Weir and Dickinson, 2012; Warren et al., 2018; Giraldo et al., 2018; Pae et al., 2024). Similarly, walking *D. melanogaster* maintain arbitrary but stable bearings relative to visual cues on their horizon (Green et al., 2019; Weisman et al., 2025). Furthermore, recent release-and-recapture experiments found that *D. melanogaster* can disperse radially at net ground speeds approaching 1 m/s, or 4 km/h (Leitch et al., 2021). This is equivalent to 1.3 million body lengths/h for a 3 mm long female. These findings raise the question of whether other *Drosophila* spp. possess similar navigation capacities to support long-distance dispersal.

This study focuses on testing the flight navigation abilities of *Drosophila suzukii* (spotted-wing drosophila), a potent, highly invasive agricultural pest whose geographic range has spread rapidly in North America and other continents over the last decade. Unlike other *Drosophila spp.,* which target already rotten fruit, female *D. suzukii* uniquely possess an enlarged, serrated ovipositor. This permits females to saw through taut fruit skin and lay eggs (Atallah et al., 2014), which develop into maggot-like larvae, ruining blueberries, raspberries, and stone fruit. Since *D. suzukii* first appeared in continental North America in 2008, it has spread rapidly and caused more than $500M of economic damage on a yearly basis (Bolda et al., 2010; Walsh et al., 2011; Tait et al., 2021). Parallel invasions, with corresponding economic damage, have occurred in South America and Europe (Asplen et al., 2015).

Despite this rapid range expansion, there is a widely held view that *D. suzukii* lacks a capacity for extended flight navigation (Kirkpatrick et al., 2018; Miller et al., 2015; Vacas et al., 2019; Rodriguez-Saona et al., 2020). For instance, an influential model posits that in the absence of local, attractive cues, *D. suzukii* flight consists of short, randomly oriented flight segments, resulting in only minimal displacement (Miller et al., 2015; Kirkpatrick et al., 2018). Consistent with this view, various release-and-recapture studies have estimated that the maximum dispersal distance for *D. suzukii* individuals is less than 90 m in a day (Kirkpatrick et al., 2018; Vacas et al., 2019; Klick et al., 2016; Rodriguez-Saona et al., 2020). Flight distance estimates obtained from tethered *D. suzukii* flying on flight mills have been similarly low, with median daily displacements of less than 30 m (Wong et al., 2018; Tran et al., 2022). Several lines of evidence, however, raise the possibility that *D. suzukii* could disperse long distances. This evidence includes the long-distance flights documented in the closely related *D. melanogaster*, as well as the reliable, annual re-emergence of *D. suzukii* in locations where seasonal climatic conditions do not permit survival (Mitsui et al., 2010; Tonina et al., 2016). This re-emergence could have resulted from progressive, long-distance reinvasion (Mitsui et al., 2010; Tonina et al., 2016). Furthermore, mark-and-recapture studies observed *D. suzukii* movement of >9 km over several weeks, though this may have been partly wind-aided (Tait et al., 2018). Notably, no prior study in *D. suzukii* has tracked continuous flight trajectories in individuals – leaving open the question of whether *D. suzukii* can maintain extended, straight flight paths. Determining the navigation capacity of *D. suzukii* is essential for understanding the species’ potential for dispersal and for planning the spatial scale of pest management and mitigation strategies.

Furthermore, the extent of sex-related differences in navigation capacity in *Drosophila* spp. is not well understood. Most tethered-flight studies of *D.suzukii* have focused on females, in part because they are larger and easier to track. A few tethered flight mill studies of *D. suzukii* have observed inconsistent differences between males and females in flight stamina, though they did not examine navigation capacity (Tran et al., 2022; Wong et al., 2018). Whereas prior studies have focused on female *D. suzukii* because they infest fruit (Wong et al., 2018), the release of sterile males is emerging as a potential control strategy (Homem et al., 2022; Yadav et al., 2023). This makes it important to understand the capacity of male *D. suzukii* to navigate and disperse.

In this study we test the navigation capacity of both male and female *D. suzukii* by continuously tracking flies’ flight headings to an overhead light source mimicking the sun. We find that both male and female *D. suzukii* maintain stable flight headings that are idiosyncratic across individuals. Our findings suggest that *D. suzukii* has an underappreciated potential for long-distance dispersal.

## Materials and Methods

### Fly rearing

We maintained *Drosophila suzukii* in a laboratory colony derived from wild-caught populations collected in and around Corvallis, Oregon, USA. To maintain robust genetic diversity in our colony, we introduced new wild-caught flies every 1-2 years and additionally used approximately 100 parent flies (50 females; 50 males) each time we generated offspring. Experimental flies were reared in bottles with cornmeal-agar-molasses diet and kept in an incubator at 20 °C under a 12 h:12 h light:dark cycle, with lights on from 08:00 to 20:00.

### Tethering

We used 2-3 days post-eclosion adult *D. suzukii* in all experiments. To prepare flies for tethering, we first cooled them to 4 °C to induce temporary immobilization. Once the flies became inactive, we positioned each fly ventral side down on a custom-built Peltier-cooled mounting platform. We then attached a steel insect pin (Living Systems Instrumentation), trimmed to 10-11 mm in length, to the anterior notum using a small drop of UV-curable adhesive (Bondic). Flies retained the ability to move their heads freely. We allowed flies to recover from cooling for at least 20 min before starting flight experiments. During this period, we gave them a small, trimmed piece of tissue (Kimwipe) to manipulate with their legs, which inhibited them from initiating flight (Giraldo et al., 2018; Pae et al., 2024; Warren et al., 2018).

### Flight Arena

We used a custom-built arena in which flies were magnetically tethered so that they could freely turn on the yaw axis but could not translate forward (Fig. 1A; Pae et al., 2024; Tammero and Dickinson, 2002; Weir and Dickinson, 2012). The flight arena consisted of a cylindrical chamber with a radius of 16.75 cm and a height of 15 cm. Each fly, attached to a steel pin, was held in place by a rare-earth rod magnet above the fly (Magcraft NSN0750, 0.125 × 1 inch) and three stacked ring magnets below the fly (Magnetic Hold, Inc.; N35EH, OD 0.75 inch × ID 0.403 inch × 0.125 inch). We positioned a low-friction jewel bearing (MT4019; Microlap) between the steel pin and the upper rod magnet, which allowed the pin to rotate freely on the yaw axis, according to the fly’s wing forces.

**Figure 1.**
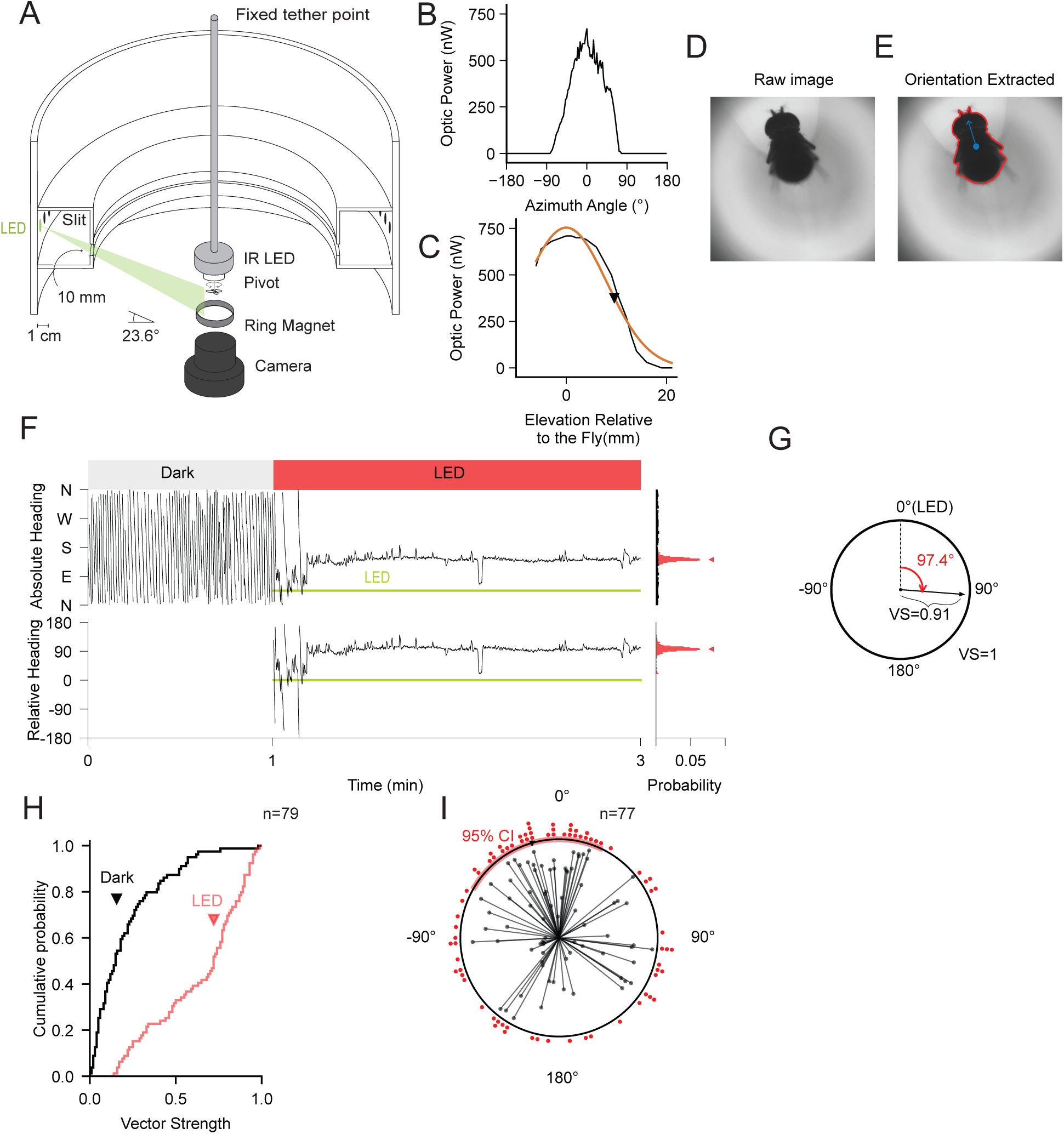
Female D. suzukii maintain stable headings relative to a discrete LED stimulus mimicking the sun. (A) Schematic of the magnetic tether arena. Flies were tethered to a steel pin, held vertically by strong magnets above and below. A low-friction bearing between the pin and overhead magnet allowed the fly to rotate freely about the yaw axis; a camera below the fly tracked body orientation voa infrared illumination. Individual green LEDs mounted at 23.6° above the fly’s horizon mimicked the sun at distinct azimuthal locations (B) Variation in power from a single LED at distinct azimuthal orientations (0°, sensor oriented toward LED; 180°, sensor oriented in opposite direction). (C) Optical power from a single LED measured at different heights. Zero corresponds to approximate location of fly. The orange curve is Gaussian fit; black marker indicates half-maximum height. (D) Example raw image. (E) Image with contour of fly’s body (red line) and heading (blue vector) extracted. (F) Representative heading traces from a female D. suzukii during initial 3 min of flight, consisting of 1 min in darkness followed by 2 min with LED activated. Top left panel shows absolute heading flown by fly during 1 min with no LED (‘Dark’) and 2 min. in which LED was activated at NE corner of arena. The bottom left panel shows fly heading relative to the LED location The green line indicates LED position in both panels. Right panels show relative probability of distinct headings during corresponding dark (black) and LED (red) periods. (G) Vector representation of the 2 min flight in (F). The mean heading is the angle of plotted vector; vector strength is magnitude of plotted vector. (H) Cumulative probability distributions of vector strength during the initial dark (black) and the first LED periods (red). Vector strength was higher during LED stimulation than during darkness, indicat-ing increased heading stability in the presence of the sun-like stimulus. Triangles indicate mean vector strength for each condition. n = 79 flights. (I) Summary of flight behavior during 2 min LED flights. Each vector corresponds to a flight, following convention of (G). The red arc indicates the 95% confidence interval of the mean heading. n = 77 flights.

We used infrared LEDs (JLCPCB; https://github.com/willdickson/magneto_ir_led_ring), not visible to *Drosophila*, to illuminate the fly’s body orientation for image acquisition during flight. To prevent ambient light from entering the arena, we enclosed it with black acrylic panels and performed experiments in a dimmed room. Individually addressable green LEDs (Adafruit NeoPixel Digital RGB LED Strip 144, WS2812 LEDs; *λ* = 515-530 nm), were used to mimic the sun. LEDs were positioned at various azimuthal positions and at 23.6° elevation above the fly’s horizon. This elevation corresponds to sun position at 8:17 am and 4:11 pm on the equinox in Corvallis, Oregon, USA, where experiments were conducted. We set LED brightness to an intensity of 30.3-38.5 mcd (2.75% of the listed 1100-1400 mcd maximum). We minimized internal reflection in the arena with a dark, non-reflective enclosure around the LEDs, through which light could only pass via a 1 cm vertical slit (Fig. 1A). Furthermore, we blackened all inner surfaces with flock paper (Edmund Optics; 54-585). To quantify the spatial extent of illumination from a single LED, we measured optical power at different heights and azimuthal orientations using a calibrated optical power meter and photodiode sensor (Thorlabs; PM400, S120C). For these measurements, we set the LED intensity 7.14 times higher than in experiments to maximize signal-to-noise ratio; we then divided measured values by the same ratio to determine optical power under experimental conditions. We fit optical power measurements with a Gaussian function,

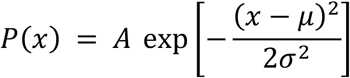

where *A* is the amplitude, *μ* is the peak position, and *σ* determines the spread.

### Data Acquisition

We controlled experiments and acquired all data with custom scripts in the ROS environment (ROS1 Noetic) on an Ubuntu 22.04 desktop computer. We recorded images [1200 x 1200 px] of flying flies at 15 Hz with a FLIR Blackfly USB 3 camera (BFS-U3-16S2M-CS) with COMPUTAR lens (MLM-3XMP). We modified a previously established computer vision package (https://github.com/willdickson/find_fly_angle) to track the fly’s body orientation continuously. We additionally saved raw images from all flies (except for 6 of 79 females, in which only partial data was saved) for post-experiment analysis and validation. In male *D. suzukii*, which are smaller and distinctly shaped from females, the body angle estimate from the previously established classifier (*find_fly_angle*) had a persistent 180° ambiguity (i.e. the directions of fly head and abdomen were not disambiguated). To resolve this ambiguity, we developed two offline classifiers. One classifier (https://github.com/warren-lab/fly-heading-correction) was a ResNet-18 convolutional neural network (CNN) (He et al., 2016), pretrained on ImageNet, and then trained on 1,675 manually labeled fly images (835 head-up, 840 head-down) from five flies, with a held-out validation set of 419 images (209 head-up, 210 head-down). We preprocessed images with Contrast Limited Adaptive Histogram Equalization (CLAHE; clip limit 2.0, 8 × 8 tile grid) to enhance local contrast, resized to 224 × 224 pixels, and normalized using ImageNet channel statistics. We replaced the final fully connected layer with a two-class head and trained 10 epochs using the Adam optimizer (learning rate 0.001, batch size 32) with cross-entropy loss. We carried out training and inference in PyTorch with torchvision and achieved 100% accuracy on the held-out validation set. We used a second, template-based classifier (https://github.com/warren-lab/flyplot) to assess and validate results from the deep learning method. For each fly, we generated a head-aligned template image by averaging 20 randomly selected images that were manually aligned. Then, we computed the pixel-wise mean squared error between the template and two image variants: the find_fly_angle alignment and an image rotated 180°. We classified body direction according to the image variant with the lower mean squared error. We used the CNN classifier to determine body orientation, except for individuals in which the percentage agreement between the two methods was lower than 98% across saved images (25 of 178). For these individuals, we manually assigned values on image frames where the two classifiers diverged.

### Experimental Protocols

Most flight trials, for both male and female *D. suzukii,* lasted 10 min. A trial consisted of an initial 1 min dark period, four consecutive 2 min periods with a single green LED activated, and then a final 1 min dark period. We permuted the LED position randomly, without replacement, across four positions (NE, NW, SE, SW) every 2 min– resulting in LED rotations of either ±90° or 180°. Following the first 10 min flight trial, we removed flies from the arena and allowed them to rest for 15 min, (Pae et al., 2024; Warren et al., 2018). During the rest period, we again provided flies with a small tissue piece to prevent flight. After the rest period, we tested the same fly again in a second trial with the same structure. The LED position sequence was independently randomized for the second trial.

In another experiment, a subset of flies flew for 17 min; here, a flight trial consisted of a 1 min initial dark period, 15 min with LED at a fixed, random position, and a final 1 min dark period. We did not perform a second flight trial in this experiment. We allowed one randomly chosen female fly to fly for 60 min continuously to a randomly chosen LED position. If a fly stopped flying during a trial, we remotely triggered a single brief air puff using an inflation needle positioned under the fly. We ended any trial in which the fly failed to resume flight or stopped repeatedly.

### Data Analysis

We analyzed all data using custom scripts in Python 3.10, using the following packages: numpy 1.21, matplotlib 3.5, PyTorch 2.12. Additional statistical analyses were performed using SciPy 1.8 and pandas 2.0.

### Heading & Vector Strength Calculations

We defined a fly’s absolute heading as the direction of the body on the yaw axis. At an absolute heading of 0°, the fly’s head was oriented north. Clockwise turns increased absolute heading. Relative heading to the LED was the azimuthal angular difference between the absolute heading and the LED position. We quantified the directional coherence of successive fly heading values using vector strength (Berens, 2009; Giraldo et al., 2018; Warren et al., 2018). We summed unit vectors corresponding to individual heading measurements and calculated the normalized magnitude of the resulting vector, which ranged between 0 (no directional coherence) and 1 (perfect coherence). We calculated heading as the direction of the resulting vector obtained by summing unit vectors that corresponded to individual relative heading measurements. For mean heading measurements, we excluded the first 10 s of flight following LED activation or LED rotation. We computed local vector strength, a dynamic, time-varying measure of heading stability by convolving a 30 s Gaussian filter window to weight unit vectors before summation of vector strength (Warren et al., 2018). We fit averaged local vector strength traces with an asymptotic model,

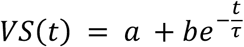

where *a* is the asymptotic vector strength, *b* is the initial offset, and *τ* is the time constant for approach to asymptote. For 15 min fixed-LED experiments, we calculated fictive displacement trajectories, assuming a constant forward flight speed of 0.5 m/s (Weir and Dickinson, 2012).

For all analyses except the descriptive vector strength distributions shown in Fig. 1H and Fig. 3E, we included only trials in which initial dark vector strength was < 0.75. High directional stability in darkness indicates that the steel pin may not have been able to rotate freely on the magnetic tether (Duistermars and Frye, 2008). This criterion excluded 2 of 79 female and 3 of 39 male single-flight experiments, as well as 6 of 59 female and 6 of 36 male paired-flight experiments.

### Quantifying behavioral responses to LED rotation

To characterize flies’ transient steering responses to LED rotations, we calculated the dynamic change in absolute heading relative to the mean over the 30 s preceding each LED rotation. We sign-inverted turning responses to −90° rotations to pool ± 90° rotations. We fit an exponential model to the mean heading trajectory following LED rotation,

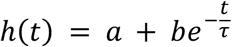

where ℎ(*t*) is the change in absolute heading, *a* is the asymptote, *b* is the initial offset, and *τ* is the time constant for approach to the asymptote.

We further compared heading differences during periods when the LED was stable and when the LED was rotated. To do this, we divided the 8 min flight period into eight 1 min intervals – and then excised the first 10 s of each interval to account for instability in heading following LED onset or rotation. We then calculated mean relative heading within each 50 s interval and computed the absolute circular difference between adjacent intervals. Adjacent interval pairs correspond to 0° LED rotations (i.e. no rotations), ±90° LED rotations, or 180° LED rotations. We compared the observed mean and distributions of pre–post heading differences across these rotation groups. Furthermore, we used these differences to examine two null hypotheses: (1) flies randomly reset their relative heading following an LED rotation and (2) flies ignored the LED rotations and behaved similarly to 0° rotations. To test whether flies randomly reset their headings after LED rotation, we randomly permuted the observed relative pre-post heading pair (n = 10,000 times) and compared the resulting null distribution of absolute circular pre-post heading differences to the observed values. To test whether flies ignored the LED rotation, we imposed pseudo-LED rotations of ±90° and 180° on flight periods in which there was no rotation.

### Statistical tests; estimation of intervals

We performed statistical analyses in Python 3.10 using custom scripts. To calculate circular means, we converted each heading measurement into a unit vector and averaged those vectors. We estimated 95% confidence intervals for circular mean headings with bootstrap resampling using 10,000 iterations.

We tested whether experiment time was correlated with fly heading by calculating the linear-circular correlation coefficient. We assessed statistical significance by randomly permuting the experiment times (while preserving fly headings) and comparing the observed correlation coefficient with the resulting null distribution. As a further test of potential time-of-day effects, we permuted time of-day labels across two time groups, while keeping group sizes the same. To test whether flies maintained a heading after a 15 min flight interruption, we randomly paired first-flight headings with second-flight headings from other flies and compared these shuffled differences to observed values. For the LED-rotation analysis, we compared the observed heading differences with null distributions obtained from the random-resetting and pseudo rotation models, as described above. All permutation tests resampled 10,000 times.

We used the Rayleigh test to evaluate the null hypothesis that distributions were uniform, and the Kruskal–Wallis test to compare absolute circular heading differences following LED rotations. When the Kruskal–Wallis test was significant, we used Dunn’s post hoc tests with Holm correction for pairwise comparisons between rotation conditions.

## Results

### Female D. suzukii maintain a stable heading to a discrete LED stimulus

We first tested whether female *D. suzukii* maintained a stable flight heading relative to a single overhead LED that mimicked the sun. We magnetically tethered flies to prevent forward translation while allowing free rotation on the yaw axis. Flies flew in a flight arena (Fig. 1A), in which the only visible light cue was one LED from a circular strip positioned 23.6° above the fly’s horizon (Fig. 1A). The azimuthal extent of LED illumination was circumscribed, with no measurable light flux from opposite directions (Fig. 1B). The vertical extent of illumination was similarly circumscribed, with a half-maximum of ∼9.8 mm above the fly (Fig. 1C). As flies flew, we imaged their body orientation at 15 fps under infrared illumination and extracted the flight heading using a previously established classifier (Fig. 1D-E). An example experiment, testing a female’s capacity to orient to a single LED, is shown in Fig. 1F. As expected, in the dark, without any available orientation cue, the fly turned erratically in an uncontrolled fashion, failing to maintain a stable heading (‘dark’, Fig. 1F). In contrast, when we activated a single green LED, the fly stabilized its heading in several seconds, then maintained a mean absolute heading of 142° for the next 2 min (97° relative to the LED; Fig. 1F, right panel). For this example, vector strength – which measures coherence of successive orientation values on a scale of 0 to 1 – was 0.91 for the LED period and 0.01 for the preceding dark interval (Fig. 1G). Across a large population of female *D. suzukii* (n = 79), we observed a similar tendency for flies to maintain a stable heading relative to a stationary LED. The mean vector strength during this initial 2 min LED period was 0.64 ± 0.03, compared to 0.22 ± 0.02 during the preceding dark period (Fig. 1H). The distribution of heading values flown relative to the LED was broad, with a circular variance of 0.36 and a 95% confidence interval of the mean that spanned 83.7° from −59.1° to 24.7° (Fig. 1I).

### Female D. suzukii make rapid, compensatory turns to maintain a stable, relative heading following rotations of a discrete sun stimulus

Female flies could have exhibited stable flight headings (cf. Figure 1) by actively maintaining a specific target azimuthal offset angle from the LED position. Alternatively, however, flies could have flown straight merely by minimizing yaw rotation, without an explicit target heading. A further possibility is that light from the LED stimulus could have illuminated other visual cues in the arena that flies used to navigate. To distinguish between these possibilities, we systematically rotated the LED position and monitored flight responses (Pae et al., 2024; Weisman et al., 2025). We reasoned that if flies actively navigated to maintain a constant offset angle from the sun stimulus, they would turn to compensate for the LED rotation, so as to maintain a stable relative heading. We randomly permuted the LED position across four cardinal orientations (NE, SE, SW, NW) every 2 min, resulting in ±90° or 180° rotations. An example experiment is shown in Figure 2A; here a female *D. suzukii* responded to three successive counterclockwise (CCW) 90° LED rotations by rapidly turning to maintain a stable relative heading. Indeed, the relative headings to the LED were much more closely distributed than the corresponding absolute headings (relative headings: −46.2°, −106.7°, −70.9°, −80.9°; absolute headings: −1.2°, −151.7°, 154.1°, 54.1°).

**Figure 2.**
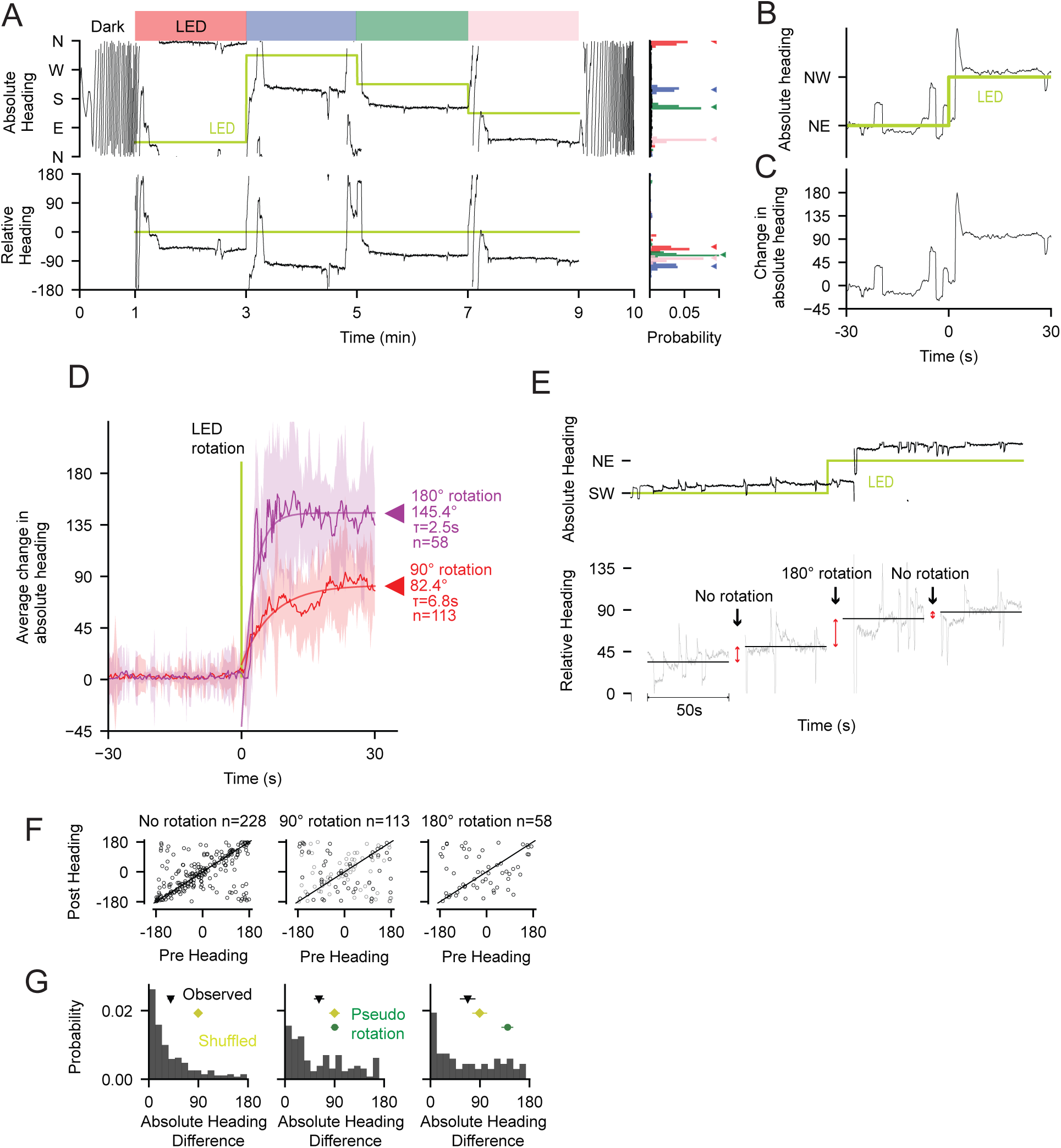
Female D. suzukii make compensatory turns to maintain stable headings following rotation of a sun stimulus. (A) Example response of a female D. suzukii to LED rotations. Top left: absolute flight heading over 10 min. The LED was off for the first and last 1 min (‘Dark’); for the remaining 8 min, we rotated the LED between four positions for 2 min each (‘LED’, all rotations were 90°). Bottom left: Relative heading flown to the LED. Right panels: relative probabilities of distinct headings. Histogram colors correspond to the left panel. (B) Example turning response (from a different fly) to a single 90° LED rotation. Top panel: Absolute heading and LED position from 30 s before to 30 s follow-ing LED rotation. (C) Change in absolute heading, calculated by subtracting the mean heading flown over 30 s prior to LED rotation. (D) Mean changes in absolute heading following LED rotation (±90° rotations, red; 180° rotations, purple). Shaded regions indicate SEM; smooth lines show fitted asymptotic models (Δ = 90°, n = 113 rotations; Δ = 180°, n = 58 rotations). (E) Example showing heading differences calculated over equivalent time periods in which the LED was stable and rotated. Top panel: Absolute flight heading (black) and LED position (green) for 2 min prior to and following a 180° rotation. Bottom panel: Relative heading for example flown over four equally spaced 50 s intervals. Mean heading differences between adjacent intervals are shown with vertical lines (corresponding to two 0° and one 180° LED rotation). (F) Pre- and post-interval relative headings for Δ = 0°, Δ = ±90°, and Δ = 180° LED rotations. (G) Distributions of absolute heading differences. Black triangles mark observed mean differences. Yellow diamonds and horizontal bars indicate the mean and central 95% interval from null model in which headings were randomly permuted. Green circles and horizontal bars indicate the mean and central 95% interval from a null model in which pseudo-rotations were imposed on Δ = 0° data, n = 228 intervals; Δ = 90°, n = 113 intervals; Δ = 180°, n = 58 intervals.

To assess the magnitude and dynamics of turning responses across a broader population of female *D. suzukii,* we aligned heading traces to the pre-rotation mean heading (Fig. 2B-C). For ±90° LED rotations, the best-fit asymptote of the mean flight response was an 82.4° turn, in the direction of LED rotation, with a time constant of 6.8 s (Fig. 2D; n = 113 rotations; 57 females). For 180° LED rotations, the best-fit asymptote was 145.4°, with a time constant of 2.5 s (Fig. 2D; n = 58 rotations; 57 females). These data indicate that female *D. suzukii,* on average, rapidly respond to rotations of the sun stimulus with compensatory steering maneuvers to maintain a stable relative heading.

To further examine the magnitude and reliability of female flies’ compensatory turns, we compared flight behavior across neighboring periods with and without LED rotation. We divided flight trials into a series of neighboring 1 min intervals (Fig. 2E). Each pair of intervals contained a 0°, ±90°, or 180° LED rotation. We calculated the mean heading difference across neighboring intervals, excluding the first 10 s to account for transient flight responses. For example, Fig. 2E shows three successive pairs of 1 min intervals (two with 0° rotation, one with 180° rotation). In this example, the mean heading differences across intervals with no LED rotation were smaller than those with rotation (16.5° and 7°, no rotation; 30.2°, 180° rotation). Similarly, across a larger dataset of female *D. suzukii*, the average heading differences for intervals with no rotation were significantly smaller than those for ±90° and 180° rotations (Fig. 2F-G; 39.7°, 61.6°, and 67.7°; n = 228, 113, 58; Kruskal–Wallis test, H = 21.67, p < 0.001; Dunn’s post hoc tests with multiple-comparison correction (no-rotation vs. 90°: p < 0.001; no-rotation vs. 180°: p = 0.002; 90° vs. 180°: p = 0.83). These results indicate that flies’ relative heading is less stable when the LED is rotated than when it is stationary.

Given there is some instability in relative heading following LED rotations (Fig. 2F), we tested the null hypothesis that flies randomly reset their headings after LED rotations. Contrary to this null hypothesis, the mean heading differences for both 90° and 180° rotations were significantly smaller than those obtained via random permutation (Fig. 2G, 90° rotation: observed mean 61.6° vs shuffled mean 89.8°, p < 0.001; 180° rotation: observed mean 67.7° vs shuffled mean 89.3°, p = 0.001). We additionally tested the null hypothesis that flies ignored the LED rotation altogether by imposing pseudo-LED rotations on baseline data. The relative heading differences following actual LED rotations were significantly lower than those calculated following pseudo-rotations (Fig. 2G, 90° rotation, observed mean difference 61.6°; pseudo-rotation mean difference 90.3°, central 95% interval: [83.2°, 97.3°], p = 0.001; 180° rotation, observed mean difference 67.7°: pseudo-rotation mean difference 140.3°, central 95% interval: [129.3°, 150.4°], p < 0.001). Together, these data indicate that female *D. suzukii* compensate for LED rotations to maintain a stable relative heading, albeit with increased heading variability. This indicates that females navigate using the azimuthal offset from a discrete sun stimulus as a target heading.

### Male D. suzukii actively maintain a stable heading to a discrete sun stimulus

We next tested whether male *D. suzukii* exhibit a similar capacity to navigate to a single LED stimulus. Because males are smaller and differently shaped than females, we developed two new classifiers to disambiguate the head and abdomen direction from saved images. One classifier used a convolutional neural network (CNN) trained via deep learning; the other measured deviations from a fly-specific template (Methods, Fig. 3A-B). The two classifiers produced highly consistent orientation estimates, with mean percentage agreement (across flies) of 99.3% (98.9% for males and 99.5% for females). There was much higher divergence between the two new classifiers and the previously established method for males than females. For males, the mean percentage agreement between the previous method and our CNN classifier was 75.4%, whereas it was 97.4% for females (Fig. 3C).

**Figure 3.**
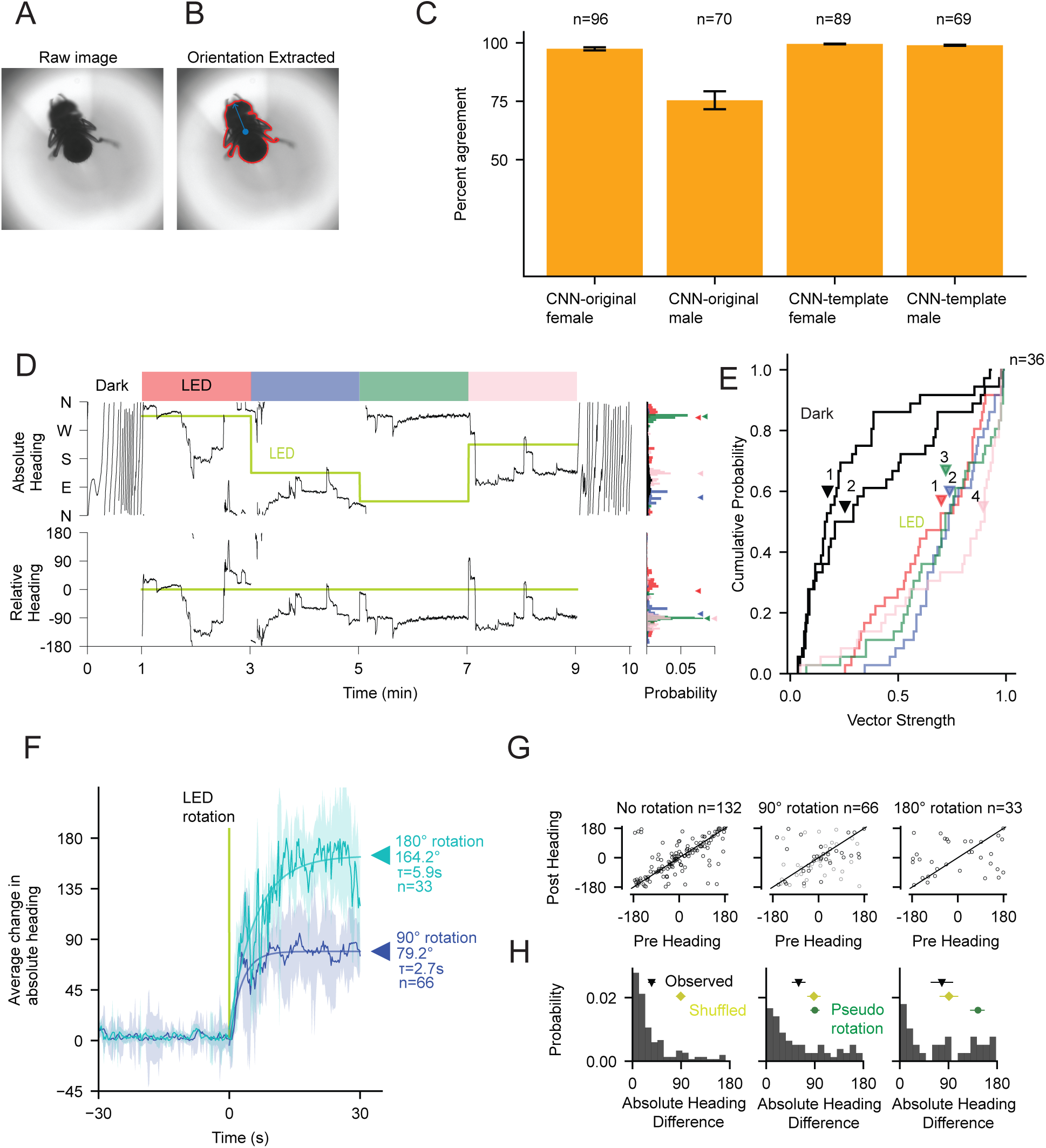
Male D. suzukii actively maintain stable headings to a sun stimulus. A) Example raw image frame used to track body orientation in a tethered flying male D. suzukii. (B) Image with body orientation extracted via a deep learning classifier. CNN confidence: 1.00, MSE ratio: 0.81. (C) Agreement among heading angle correction methods. Bars show the percentage of frames for which the compared methods produced matching head-abdomen assignments; error bars indicate SEM across flies. (D) Example flight response of a male D. suzukii in response to LED rotations. Top left panel: Absolute flight heading during 1 min dark period, followed by four 2 min periods with LED at four distinct directions (NW, SE, NE, and SW). Bottom left panel: Flight heading relative to LED direction, shown from −180° to 180°. Right panels: probability distributions of absolute and relative heading during the corresponding dark and LED periods. (E) Cumulative distributions of vector strength during two dark and four LED periods. (F) Mean change in absolute heading following LED rotations in male flies. Absolute headings were calculated relative to the mean value during the 30 s pre-ro-tation period. Responses are shown for ±90° (pooled via sign inversion) and 180° LED rotations; shaded regions indicate SEM, and smooth curves show exponential fits. Δ = 90°, n = 66 rotations; Δ = 180°, n = 33 rotations. (G) Pre- and post-interval relative headings calculated over neighboring 50 s intervals, as illustrated in Fig. 2E. Scatter plots with pre-and post-rotation mean relative headings. (H) Distributions of absolute pre-post heading differences. Black triangles indicate observed mean differences. Yellow diamonds and horizontal bars indicate the mean and central 95% interval from a shuffied null model in which post-interval headings were randomly reassigned. Green circles and horizontal bars indicate the mean and central 95% interval from a pseudo-rotation null model calculated during Δ = 0° intervals. Δ = 0°, n = 132 intervals; Δ = 90°, n = 66 intervals; Δ = 180°, n = 33 intervals.

We tested males’ flight navigation capacity with the same experimental approach used for females. An example flight, in which the location of a discrete sun stimulus was rotated every 2 min, is shown in Fig. 3D. Here the male fly responded to the LED rotations (of 180°, −90°, and 180°) with turns that maintained a stable relative heading (relative headings were −4.8°, −77.5°, −91.3°, −92.1°). Across a larger population, the mean vector strength for each of the four flight segments closely matched values observed in females (Fig. 3E, males: 0.66, 0.74, 0.71, 0.74; females: 0.65, 0.72, 0.76, 0.73). Similarly, the magnitude and temporal dynamics of male flight responses to LED rotations closely matched females. The fitted asymptote of the mean turning response was 79.2° for ± 90° rotations and 164.2° for 180° rotations (Fig. 3F, n = 66 ± 90° rotations; 33 180° rotations. 33 individuals). The time constants for 90° and 180° rotations were 2.7 s and 5.9 s, respectively, compared with 6.8 s and 2.5 s for females. As with females, the mean heading differences between neighboring 1 min periods were significantly larger following 90° and 180° LED rotations than after no rotation (Fig. 3G-H; no rotation: 35.4°, ±90° rotation: 60.0°, 180° rotation: 77.8°, respectively; n = 132, 66, 33; Kruskal–Wallis test, H = 18.20, p < 0.001; Dunn’s post hoc tests with multiple-comparison correction (no-rotation vs. 90°: p = 0.003; no-rotation vs. 180°: p = 0.001; 90° vs. 180°: p = 0.34). For both ±90° and 180° LED rotations, the heading differences were significantly smaller than values obtained by applying pseudo-LED rotations on baseline data (Fig. 3H, 90°: pseudo-rotation mean 90.0°, central 95% interval: [81.7°, 98.7°], p < 0.001; 180°: pseudo-rotation mean 144.6°, central 95% interval: [130.1°, 156.9°], p < 0.001) – further indicating male flies responded to the LED rotation to maintain a stable heading. For ±90° LED rotations, the observed heading difference of 60.0° was significantly lower than values obtained by randomly permuting flight segments (n = 66, p < 0.001). However, the heading difference for 180° LED rotations, 77.8°, was not significantly smaller than differences obtained by permutation (n = 33, p = 0.067). In summary, the magnitude and dynamics of steering responses, and the clear differences from periods with no LED rotation, indicate male *D. suzukii* actively navigate using a discrete sun stimulus as a reference cue – though for 180° LED rotations, we cannot rule out the possibility of random heading resetting.

### Male and female flight headings are near uniformly distributed and not explained by time of day

The ability of *D. suzukii* to maintain a stable course relative to a rotating sun stimulus (Figs. 2 and 3) raises the question of what flight headings individual flies choose. To address this question, we calculated the net overall heading flown, relative to the LED, using flight data from all four 2 min flight segments (Fig. 4A). The distribution of relative headings, calculated across all flies, was broad and nearly uniform. The 95% confidence interval for the mean heading spanned 314.8° (−162° to 152.8°), leaving only a 45.2° region outside the interval (Fig. 4B, n = 90). Confidence intervals for male and female *D. suzukii,* when calculated separately, were similarly broad (males: −144.2° to 119.8°; females: −172.9° to 171.2°). For the combined data, as well as males and females considered separately, we could not reject the null hypothesis of a uniform distribution (Rayleigh test; combined: p = 0.61, males: p = 0.47, females: p = 0.62).

**Figure 4.**
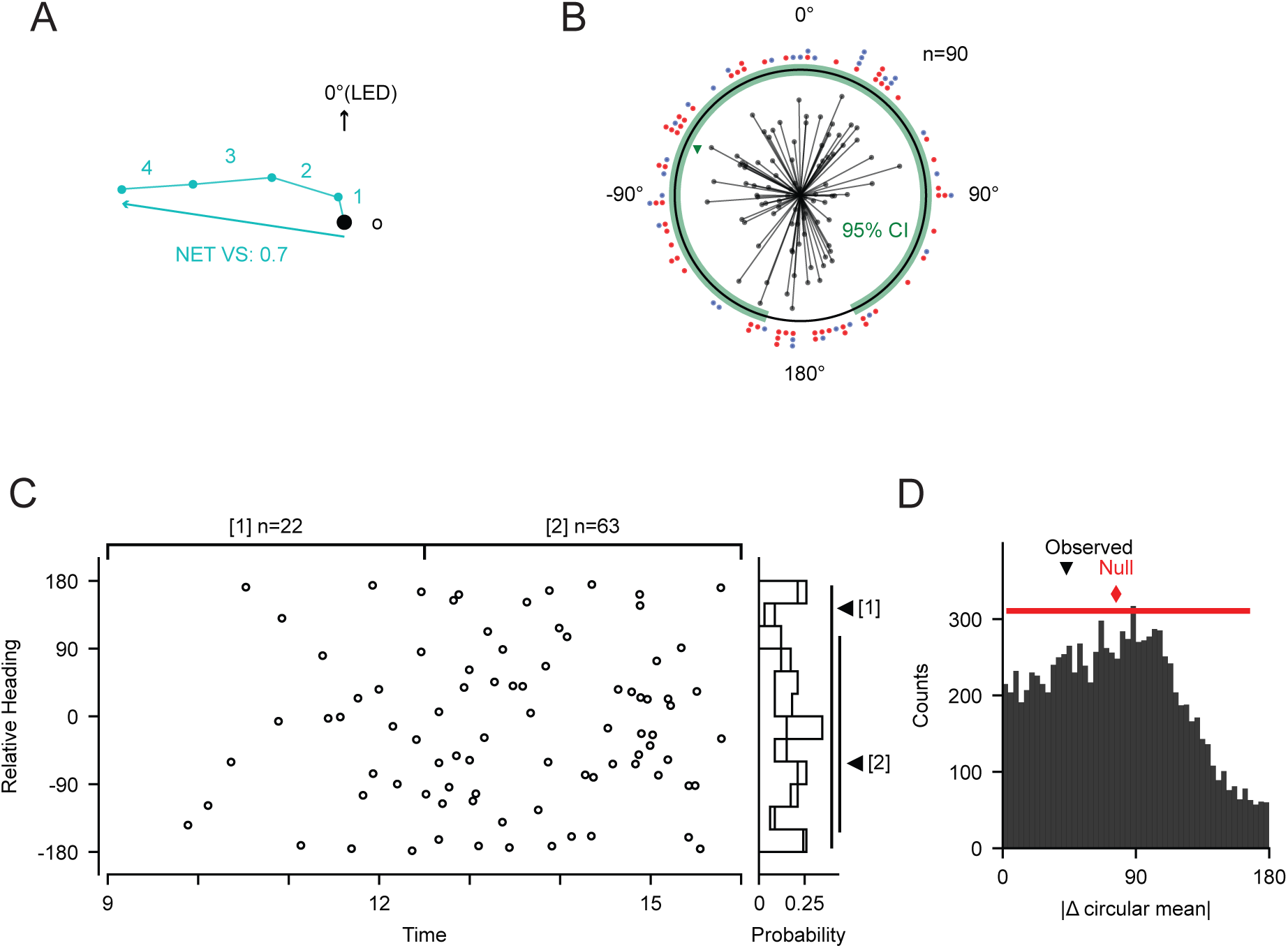
D. suzukii Hy at stable but idiosyncratic headings relative to a sun stimulus. (A) Schematic showing how we calculated relative headings from the four 2 min LED periods to estimate each fly’s net heading preference across the full 8 min LED period. (B) Distribution of net headings flown by male and female flies across 8 min flights. Each radial line represents the mean relative heading and vector strength of an individual flight during the full 8 min LED period. Red and blue dots indicate the distributions of female and male mean headings, respectively. The green triangle marks the overall circular mean, and the green arc indicates the bootstrapped 95% confidence interval, which spanned from −162° to 152.8° for the population mean heading. n = 90. (C) Relationship between trial time and net relative heading during LED stimulation. Each point represents one flight plotted against time of day. Data were divided into earlier and later time groups, indicated by brackets [1]: n = 22 and [2]: n = 63. Right, probability distributions of mean relative headings for each time group; triangles indicate circular mean heading, and vertical bars denote the bootstrapped 95% confidence intervals. Group sizes are indicated above the plot. (D) Permutation test for the time-of-day effect shown in (C). The histogram shows the null distribution of absolute circular differences in circular mean relative heading between the two time groups after random permutation of group labels. The black triangle indicates the observed difference, and the red triangle indicates the mean of the permutation null distribution.

One factor that could have potentially explained variation in flight heading is the time of day that experiments took place. However, we found no significant linear-circular correlation between time of day and net relative heading (overall: n = 85, r = 0.143, p = 0.433; female: n = 56, r = 0.196, p = 0.350; male: n = 29, r = 0.299, p = 0.303; Fig. 4C). We also compared mean heading between two equal-duration time windows (9:00-12:30, n = 22; 12:30-16:00, n = 63). The observed difference in mean heading between these time windows was not significantly different from the range of differences obtained by randomly reassigning data across the two periods (p = 0.73; Fig. 4D).

### *D. suzukii* maintain their initial heading preference following a 15 minute interruption to flight

The data presented so far raise the question of whether individual *D. suzukii* maintain their particular heading preferences following the cessation of flight. Recent studies have shown that flying *D. melanogaster* maintains individual heading preferences for tens of minutes (Giraldo et al., 2018; Pae et al., 2024) and that walking *D. melanogaster* do so for days or even weeks (Weisman et al., 2025). To test the stability of *D. suzukii* heading preference, we kept flies from flying for 15 min after the termination of the first flight and then initiated a second 10 min flight, using the same experimental paradigm. An example experiment, with a *D. suzukii* female, is shown in Fig. 5A. In the first 10 min flight, the fly flew at a net relative heading of −78.13° (Fig. 5A; individual segments −70.0°, −52.3°, 109.5°, and −106.9°). In the second flight, the fly flew at a nearly identical net heading, of −73.18° (Fig. 5A; individual LED headings of −46.0°, −104.9°, −70.6°, −81.1°).

**Figure 5.**
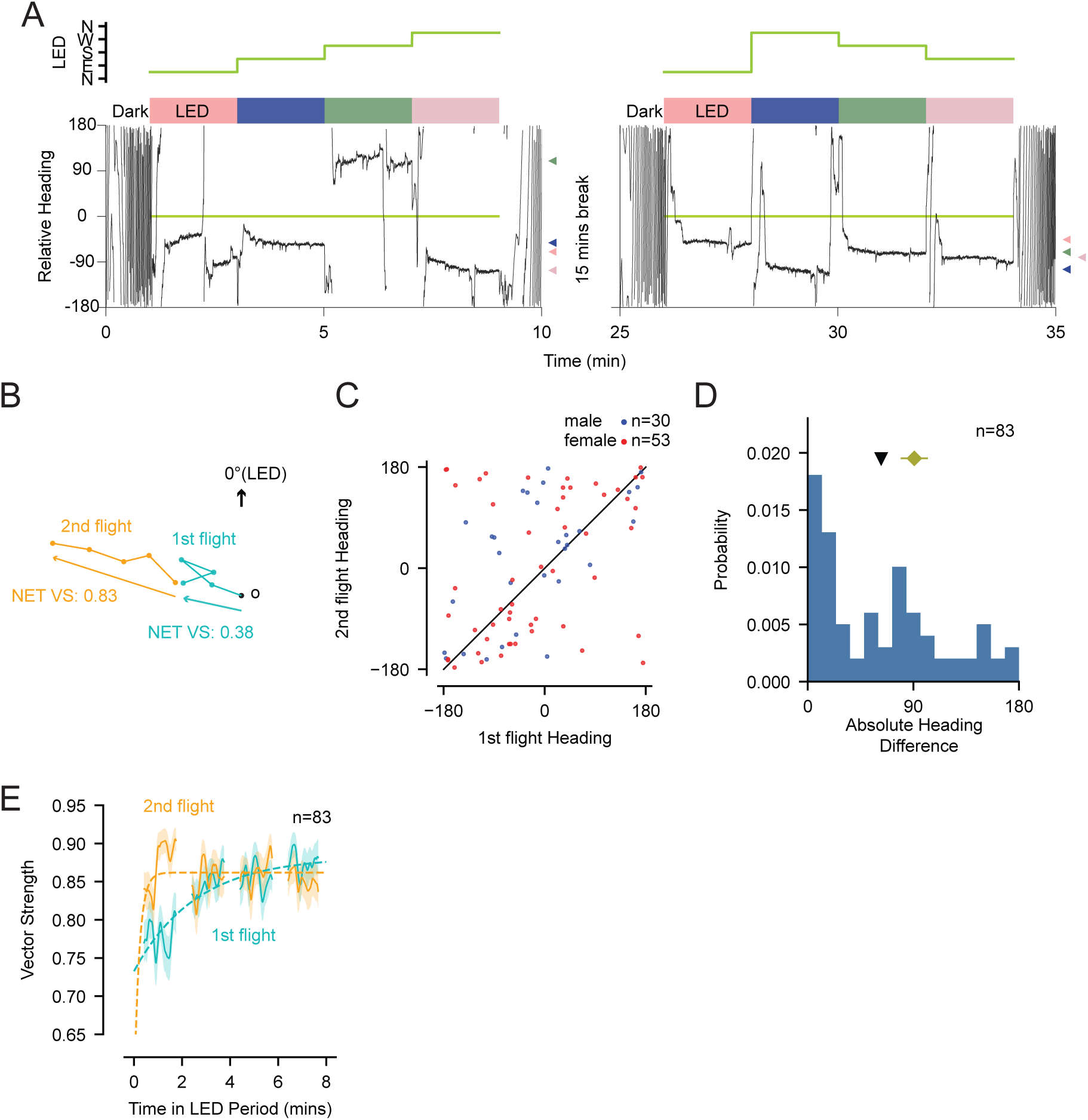
Flies maintain their initial heading following a 15 min Hight interruption, with more rapid heading stabi-lization in the second Hight. (A) Example flights flown by female D. suzukii over two flight trials separated by a 15 min interruption. The black trace shows relative heading to the LED direction. Colored bars indicate LED periods, dark periods are labeled above the trace, and LED positions are shown in compass coordinates. LED positions were presented in randomized order in each trial. (B) Vector summary of the example paired-flight trial shown in (A), illustrating the similarity between the fly’s net relative heading in the first and second flights. (C) Relationship between relative headings in the first and second flight trials. Red and blue points indicating female and male flies, respectively. Mean relative heading was calculated across the LED periods within each flight. The diagonal line indicates identical mean relative heading across the two trials. (D) Distribution of absolute heading differences between the first and second flight trials. The black triangle indicates the observed mean difference. The yellow diamond and horizontal bar indicate the mean and central 95% interval of a shuffied null distribution generated by randomly reassigning first-second flight pairs. n = 83 flies. (E) Time course of mean local vector strength during the LED periods in the first and second flight trials. Solid lines show mean values, shaded regions indicate SEM, and dashed curves show exponential fits. n = 83 flies.

Across 83 *D. suzukii* (30 males and 53 females), there was a significant tendency for flies to maintain their initial heading preference in a second flight (paired data; Fig. 5C). The mean absolute heading difference between the two flights was 62.5°, significantly smaller than values obtained by random reshuffling of flight headings (null mean 90.7°, p < 0.001) (Fig. 5D). There was an effect on both sexes when analyzed separately. Male flies showed a mean absolute first-second flight difference of 72.6°, compared with a shuffled null mean of 92.8° (n = 30, p = 0.016), whereas female flies showed a mean absolute first-second flight difference of 56.8°, compared with a shuffled null mean of 88.2° (n = 53, p < 0.001). These data indicate that flies maintained a heading preference for at least 15 min after flight cessation. We compared the temporal evolution of heading stability in the first and second flight with a time-varying estimate of vector strength. We found that the flight heading stabilized over several minutes in the initial 10 min flight (Fig. 5E, *τ* = 154 s) but within 15 s in the second flight (Fig. 5E, *τ* = 12.5 s). Taken together, these data suggest that *D. suzukii* retain their individual heading preferences following a 15 min interruption to flight; furthermore, flies rapidly stabilize their flight heading when initiating a second flight but not an initial one.

### D. suzukii exhibit stable headings over extended flights at a single LED position

*D. suzukii’s* compensatory responses to LED rotations suggest that flies would maintain an extended, stable absolute heading if the sun stimulus remained stationary. This is relevant to field dispersal under the open sky – where the sun position remains stable over a time scale of tens of minutes. We conducted a series of 15 min flights with female *D. suzukii* in which the LED remained stationary at a randomly chosen location. As predicted, we found that flies maintained highly stable headings. An example is shown in Fig. 6A. Here, the fly maintained a relative heading of 50.8° – with a vector strength of 0.79 over the 15 min flight. Assuming a constant ground flight speed of 0.5 m/s, this would correspond to a 355 m displacement. Across a broader population (n = 18 females, 3 males), we observed that flies similarly maintained stable headings over a broad range of heading values (mean vector strength 0.61 ± 0.04), corresponding to fictive displacements of 275 ± 18m in 15 min at flight speeds of 0.5 m/s (Fig. 6B). Females’ mean vector strength was 0.59 ± 0.04, whereas males’ was 0.75 ± 0.09. In one randomly selected female, we extended the fixed-LED experiment to 60 min. This fly maintained a stable relative heading of −45.3° with a vector strength of 0.91, which corresponds to 1.64 km at a flight speed of 0.5 m/s (Fig. 6C). Together, these data indicate that *D. suzukii* maintain stable, arbitrary headings to a sun stimulus for extended flight periods, extending up to an hour.

**Figure 6.**
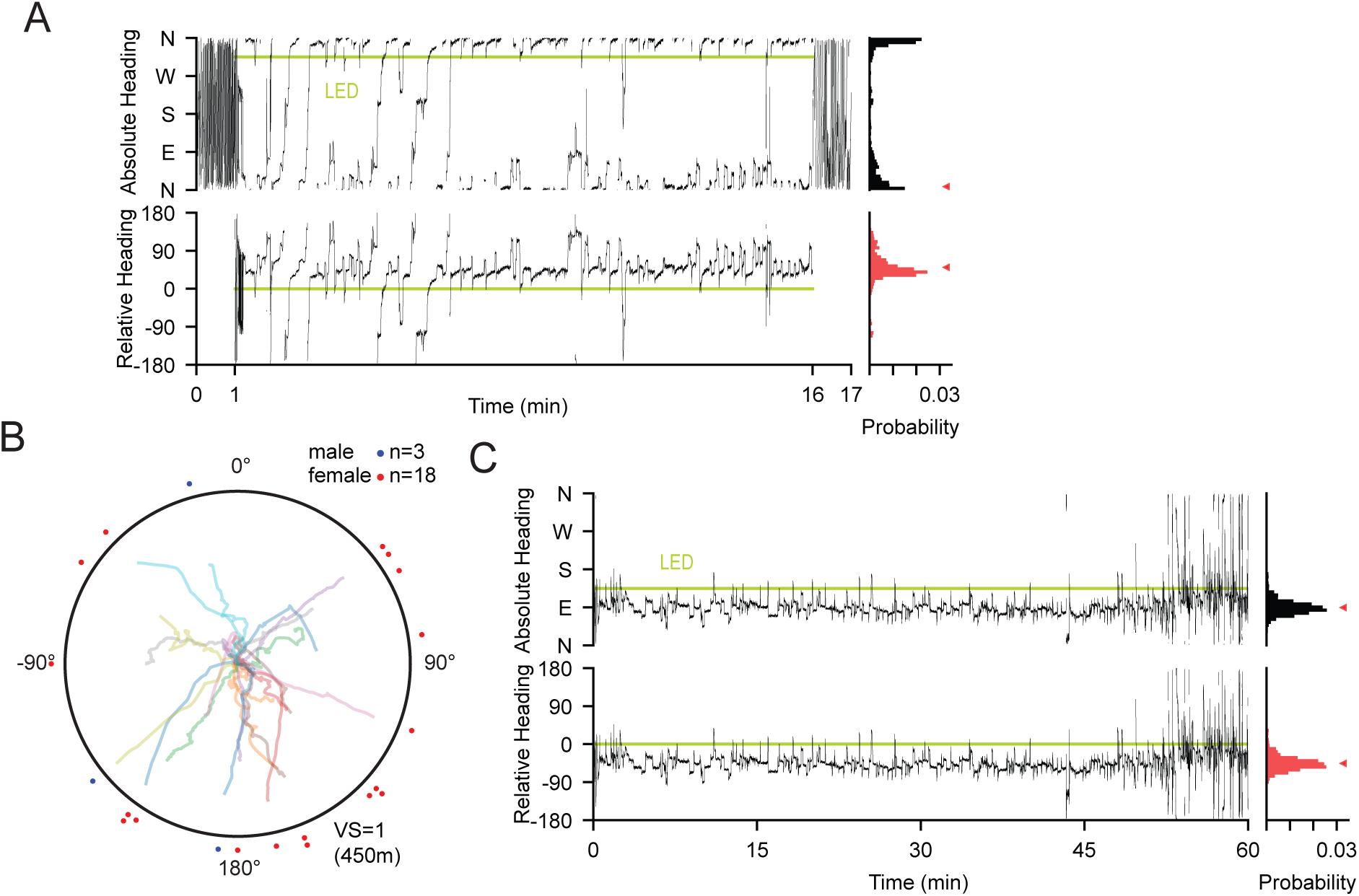
D. suzukii maintain stable headings relative to a stationary sun stimulus over extended Hight periods. (A) Representative heading behavior of a tethered female D. suzukii during a 17 min flight consisting of 1 min dark, 15 min with stationary LED at NW position, and a final 1 min dark period. Top, absolute flight heading displayed on a compass scale, with cardinal directions labeled as N, E, S, W. Bottom, heading relative to the LED direction, shown from −180° to 180°. The horizontal light-green line indicates the LED position. Right, probability distributions of absolute and relative heading during LED period. (B) Reconstructed fictive flight trajectories from 21 individual 15 min fixed-LED flights, including 18 female and 3 male flights. Trajectories were calculated by integrating measured heading over time while assuming a constant forward flight speed of 0.5 m/s. The outer circle represents the displacement expected for a perfectly straight 15 min flight, equivalent to vector strength = 1; under the assumed flight speed, this corresponds to 450 m. Points around the perimeter indicate individual net relative headings. (C) Example 60 min flight with the LED fixed at one azimuthal position (SE). Absolute and relative heading traces are shown over 60 min, with the LED position indicated by the horizontal light-green line. Right, probability distributions of absolute and relative heading.

## Discussion

In this study, we tested how flying *D. suzukii* navigate relative to a discrete overhead LED stimulus that mimicked the sun. Both male and female *D. suzukii* flew at stable, straight headings, which varied idiosyncratically across individuals. Across the population, flies’ flight headings relative to the sun stimulus were broadly distributed. Flies’ individual heading preferences persisted for at least 15 min following the cessation of an initial flight. Our findings suggest that *D. suzukii* have an underappreciated capacity for stable flight navigation, which could enable rapid, radial dispersal.

Our findings challenge a commonly held assumption that in the absence of local, attractive cues, *D. suzukii* adopt short flight segments that lack directional coherence (Kirkpatrick et al., 2018; Miller et al., 2015). Unlike prior tethered flight mill studies (Tran et al., 2022; Wong et al., 2018), we recorded continuous flight trajectories while flies were actively navigating. The mean vector strength in 15 min flights with a stationary sun stimulus was 0.61, similar to values (0.69 and 0.72, for males and females respectively) in our larger dataset of 2 min flight segments. How far flies could disperse with this degree of heading straightness depends on their flight speed. Tethered flight mill experiments in *D. suzukii* have recorded mean flight speeds of 0.18-0.30 m/s, with maximum values exceeding 0.5 m/s (Tran et al., 2022; Wong et al., 2018). However, tethered flight studies likely significantly underestimate the speed flown in free flight (Asplen, 2018; Heinrich, 1971; Taylor et al., 2010),. Indeed, in wind tunnel experiments, *D. suzukii* routinely fly upwind against a 0.3 m/s laminar flow (Duménil et al., 2025; Hemer et al., 2025; Rehermann et al., 2022). In the smaller *D. melanogaster,* instantaneous flight speeds can reach 1.6 m/s, and flies can cover 1 km at net speeds of 1 m/s, implying faster instantaneous velocities (Leitch et al., 2021; Ray et al., 2016). We based our fictive displacement estimates (Fig. 6B) on a 0.5 m/s flight speed, as done previously for *D. melanogaster* (Weir and Dickinson, 2012). Using this flight speed, we estimated a mean displacement of 275 m for *D. suzukii* in 15 min, and a maximum of 405 m (VS = 0.9). This suggests that *D. suzukii* could cover a km in an hour, as observed in *D. melanogaster* (Leitch et al., 2021). Indeed, in this study, a single *D. suzukii* flew 60 min without feeding with a vector strength of 0.91, corresponding to an estimated dispersal distance of 1.64 km (Fig. 6C).

These estimated dispersal distances for *D. suzukii* are ∼10 times larger than values measured in release-and-recapture experiments (Kirkpatrick et al., 2018; Rodriguez-Saona et al., 2020; Vacas et al., 2019). One possible explanation for this discrepancy is that the release-and-recapture experiments underestimated flies’ dispersal speed by only checking the baited traps daily. Indeed, our results suggest that some flies might have reached 100 m traps in tens of minutes rather than in a day. Another potential explanation is that flies’ impulse to disperse (and thereby use their navigation capacity) depends on specific environmental and/or fly-intrinsic cues such as hunger (Johnson, 1969). Indeed, a recent study documenting rapid, radial dispersal in *D. melanogaster* conducted release experiments in barren, desert locations with protein-deprived flies (Leitch et al., 2021). Future release experiments in *D. suzukii* could test dispersal speed directly, using a computerized camera trap approach that timestamps arrival time (Leitch et al., 2021). Furthermore, future experiments could test how environmental conditions (e.g. site conditions, wind, temperature) and intrinsic factors such as hunger influence dispersal.

The stable, idiosyncratic flight headings flown by *D. suzukii* are broadly similar to navigation behaviors recently documented in *D. melanogaster*. In many cases, flies exhibit menotaxis, by maintaining an arbitrary but stable angular offset from a reference cue (Giraldo et al., 2018; Green et al., 2019; Pae et al., 2024; Sullivan et al., 2019; Warren et al., 2018; Warren et al., 2019; Weisman et al., 2025). This includes flight headings to various celestial cues, including the overhead pattern of polarized light, sun stimuli, and sky-wide intensity cues (Giraldo et al., 2018; Sullivan et al., 2019; Warren et al., 2018; Warren et al., 2019) and walking responses to vertical cues on flies’ horizon (Green et al., 2019; Weisman et al., 2025). Together, these results suggest menotaxis could be a general navigation strategy for insects dispersing from a central location without a definite target (Warren et al., 2019). Indeed, dung beetles, when rolling dung balls away from a starting location, exhibit similarly stable and idiosyncratic headings (Dacke and el Jundi, 2018). Menotaxis at unpredictable headings may ensure that some subset of a larger population reaches refuge, food, or potential mates (Warren et al., 2019). Indeed, environmental filtering of insects that disperse radially could explain some instances of apparent directional migration in *Drosophila* spp. and other Dipterans (Hawkes et al., 2025).

Our particular finding that *D. suzukii* makes rapid, compensatory turns following rotations of a discrete sun stimulus contrasts with a recent study in *D. melanogaster* (Pae et al., 2024). That study found that flies did not maintain stable headings following rotations of a single LED; however, they did make compensatory adaptive heading adjustments following rotations of a dual sun/anti-sun stimulus, with two LEDs offset across the artificial sky by 180° (Pae et al., 2024). Our results in this study suggest that *Drosophila suzukii* can navigate with a partial view of the sky, if flies are able to see the sun. The ability to navigate to a partial sky view is essential for robust navigation (Brines and Gould, 1982; Gould, 1995; Gould, 1998). The distinct turning responses in these studies could reflect fundamental differences in sun navigation between species (i.e. *D. suzukii* might be innately more capable at navigating to an isolated sun stimulus and partial view of the sky). An alternative, not mutually exclusive explanation, is that the discrepancy in turning responses reflects subtle differences in experimental conditions, such as LED brightness. Notably, walking *D. melanogaster* turn to maintain stable headings following rotations of a single vertical cue on their horizon (Weisman et al., 2025); tethered, flying monarch butterflies perform similar compensatory turns following rotations of a discrete sun stimulus (Franzke et al., 2022). In summary, our results indicate *D. suzukii* actively navigate using a discrete sun stimulus as a reference cue; the extent to which *D. melanogaster* can do the same deserves further testing.

Whereas most prior studies of tethered flight in *D. melanogaster* and *D. suzukii* have focused on females, we studied navigation in both male and female *D. suzukii*. Females’ larger size permits more straightforward tethering and tracking; however, here we showed that male *D. suzukii* body orientation can be disambiguated via a deep learning image classifier (Fig. 3). We found that male and female *D. suzukii* have a similarly robust capacity for sun navigation (Fig. 2, 3). The bulk release of sterile male *D. suzukii* is emerging as a potential technique to manage wild populations (Homem et al., 2022; Yadav et al., 2023). Our results suggest that male *D. suzukii* possess the behavioral capacity for long-distance, radial dispersal, supporting these sterile release strategies.

The interindividual variation we observed in flight headings raises the question of how flies choose a target direction. In *D. melanogaster*, numerous behavioral phenotypes are highly variable across individuals yet stable for a particular fly (Bivort, 2025; Buchanan et al., 2015; Godesberg et al., 2024; Mathejczyk et al., 2026), raising the possibility the behaviors are genetically controlled or fixed immutably during development (Bivort, 2025; Buchanan et al., 2015; Linneweber et al., 2020). Here we observed that flies’ flight headings stabilized over several minutes in the first of two paired flights but almost immediately in a second flight (Fig. 5E). This supports the view that flight heading preference is not hard-wired but instead coheres around a random value gradually (Warren et al., 2018). Consistent with this view, flying *D. melanogaster* do not retain a heading preference after a 6 hr interruption to flight (Giraldo et al., 2018; Pae et al., 2024). In contrast, in walking *D. melanogaster*, individual heading preferences may persist for days or weeks (Weisman et al., 2025). An open question is whether, and in what contexts, *Drosophila* spp. can directly select or influence their flight heading relative to celestial reference cues. This ability to select and maintain specific azimuthal offset angles may underlie the directed, long-distance migration performed by monarch butterflies and other renowned navigators (Kendzel et al., 2026).

In summary, our study indicates that both *D. suzukii* males and females have an underappreciated capacity to fly at extended, straight headings relative to an overhead sun stimulus. Under plausible assumptions about forward flight speed, this navigation capacity could support displacement over 200 m in 15 min and therefore km-scale dispersal in longer flights. Furthermore, the capacity of *D. suzukii* to navigate could underlie the reliable, annual re-establishment of *D. suzukii* in seasonally inhospitable locations (Mitsui et al., 2010; Tonina et al., 2016). Our results point to the necessity of managing dipteran pests species such as *D. suzukii* at an area-wide scale (i.e. 10-100 km), given their potential to disperse long distances.

## Acknowledgements

We thank Michael Dickinson, Will Dickson, Ysabel Giraldo, and Francesca Ponce for technical advice. We thank Jana Lee and Matoska Silva for comments on a draft version of this paper. We received funding from the USDA-AFRI NIFA and USDA-AFRI BRAG programs.

## Notes

### Competing Interest Statement

The authors have declared no competing interest.

### Summary of Updates

This version has been updated with addition of line numbers, tighter line spacing for easier reading, and minor text changes for clarity.

